# Structural insights and mechanistic inhibition of *Plasmodium falciparum* dihydroorotate dehydrogenase by newly identified chemotypes

**DOI:** 10.64898/2026.10.25.761520

**Authors:** Japheth Kibet Ng’etich, Ryuta Ishii, Hina Yuasa, Ryo Hatanaka, Aiko Hattori, Jun-ichi Kishikawa, Endah Dwi Hartuti, Normalita Eka Pravitasari, Kiyoshi Kita, Yusuke Tateno, Teruhisa Kato, Takaya Sakura, Tomoo Shiba, Daniel Ken Inaoka

## Abstract

Resistance to frontline antimalarial drugs continues to threaten malaria control, highlighting the need for new chemotypes. *Plasmodium falciparum* dihydroorotate dehydrogenase (PfDHODH) catalyzes a key step in *de novo* pyrimidine biosynthesis and remains a validated antimalarial target. Although the PfDHODH inhibitor DSM265 advanced to clinical development, resistance-associated mutations highlight the need for structurally distinct inhibitors. Here, we report the discovery and structural characterization of PfDHODH inhibitors identified through target-based high-throughput screening of 432,592 compounds. Primary hits were evaluated by dose-response assays against asexual blood-stage parasites, mammalian mitochondrial toxicity assays, and HepG2 cytotoxicity profiling, yielding 348 hits. Seven representative compounds were prioritized based on efficacy against PfDHODH and orthologs from *P. vivax* and *P. berghei*, and their X-ray co-crystal structures revealed distinct binding modes within the active site. Ferulenol exhibited sub-micromolar inhibition of recombinant PfDHODH (*r*PfDHODH) and *r*PvDHODH, together with potent blood-stage activity, with its prenylated tail contributing additional hydrophobic interactions. Compound 2 showed the highest potency across the tested DHODH enzymes but reduced selectivity for PfDHODH, potentially due to its distinct binding orientation and furylmethyl substituent. Comparative structural analyses with DSM265 identified binding-site features that may guide the design of next-generation PfDHODH inhibitors with improved potency and selectivity.

## 1. Introduction

Malaria remains a major public health challenge, with increasing cases and deaths following the COVID-19 pandemic (Huang et al., 2025). Artemisinin-based combination therapies (ACTs) remain the standard treatment for uncomplicated malaria and have substantially reduced disease burden (Grueninger and Hamed, 2013; Laktabai et al., 2024). However, the spread of drug-resistant *Plasmodium* strains, including the emergence of partial artemisinin resistance in Africa, threatens recent progress in malaria control and elimination (Pranty et al., 2024; Björkman et al., 2024; Ward et al., 2022; Watson et al., 2022; Dembele et al., 2017; Phillips et al., 2015). These trends underscore the need for new antimalarial agents with distinct mechanisms of action (Dziwornu et al., 2024). Progress toward this goal also depends on prioritizing parasite pathways with validated druggability, as the discovery of new *Plasmodium* targets remains constrained by the limited capacity of current genetic tools to systematically interrogate essential genes (Moon et al., 2013; Raphemot et al., 2016).

The *Plasmodium* mitochondrial electron transport chain (MtETC) is a validated antimalarial target and comprises several enzymes involved in electron transfer, including dihydroorotate dehydrogenase (DHODH), succinate dehydrogenase, malate quinone oxidoreductase, and type II NADH dehydrogenase (Espino-Sanchez et al., 2023; García-Guerrero et al., 2024; Vaidya et al., 2008; Sheokand et al., 2024). DHODH is a flavin mononucleotide (FMN)-dependent enzyme that catalyzes the oxidation of dihydroorotate to orotate during *de novo* pyrimidine biosynthesis. The reaction proceeds through reduction of FMN by dihydroorotate followed by FMN reoxidation by coenzyme Q. Because *Plasmodium* lacks a functional pyrimidine salvage pathway, the parasite is dependent on *de novo* pyrimidine biosynthesis, making DHODH an attractive therapeutic target (Kokkonda et al., 2020; Pontikos et al., 2022; Singh et al., 2017).

DSM265, identified through target-based drug discovery, was the first DHODH inhibitor to advance to clinical trials and demonstrated activity against both blood-and liver-stage *Plasmodium* parasites (Phillips et al., 2015). However, its clinical development was limited by toxicity and the emergence of resistance. Recrudescent parasites from treated patients harbored DHODH substitutions, including C276Y, C276F, and G181S, as well as evidence of *dhodh* gene amplification, highlighting the potential for target-based resistance. These findings further validated DHODH as an antimalarial target while emphasizing the need for chemically diverse inhibitors with improved potency, selectivity, and resistance profiles (Palmer et al., 2021; White et al., 2019).

We conducted a high-throughput screen of a 432,592-compound library from Shionogi & Co., Ltd., against *r*PfDHODH and characterized selected hits for antiplasmodial activity and target selectivity. Seven inhibitors yielded *r*PfDHODH co-crystal structures, revealing distinct binding modes within the active site. Their activities were further evaluated against DHODH orthologs from *P. vivax* (*r*PvDHODH) and *P. berghei* (*r*PbDHODH), as well as human DHODH, providing insights into species selectivity. Most compounds showed pronounced selectivity for parasite DHODH over the human enzyme and lacked detectable inhibition of mammalian mitochondrial respiratory complexes. Together, these findings expand the chemical and structural diversity of PfDHODH inhibitors and provide a framework for the development of next-generation antimalarial agents and potential partner drugs for combination therapies.

## 2. Results

### 2.1 Identification of PfDHODH Inhibitors by High-Throughput Screening

To identify novel *r*PfDHODH inhibitors, we screened a library of more than 400,000 compounds from Shionogi & Co., Ltd., using a series of assays (Fig. 1a). Before the full-scale high-throughput screening campaign, a pilot study using a smaller compound library yielded a promising hit rate, supporting the feasibility of the screening strategy. We subsequently assessed *r*PfDHODH inhibition in a single-dose assay at a final compound concentration of 3.1 µM. The primary screen was successful, with a Z′ factor of 0.89 ± 0.04, S/B ratio of 5.2, and hit rate of 1.4% (≥40% inhibition). The average IC_50_ value of the reference inhibitor DSM265 against *r*PfDHODH across screening plates was 4.2 nM, consistent with previously reported values (Phillips et al., 2016) (Fig. 1a and Fig. S2a-c). The subsequent dose–response assay for *r*Pf*DHODH* also demonstrated excellent assay robustness, with a Z′ factor of 0.90 ± 0.02, an S/B ratio of 5.2, and an average DSM265 IC_50_ of 3.85 nM. These results yielded a hit-confirmation rate of 90.7% at the ≥40% inhibition threshold (Fig. S2d–f). Of the confirmed hits, 362 compounds showed no clear inhibition of *r*HsDHODH, suggesting they were parasite-specific inhibitors (Fig. S2g). We selected compounds exhibiting high selectivity against *r*PfDHODH (*r*PfDHODH IC_50_ ≤10 µM and *r*PfDHODH/*r*HsDHODH IC_50_ ratio >3) or high activity with less selectivity (*r*PfDHODH IC_50_ ≤1 µM and IC_50_ ratio of 1/3 < *r*PfDHODH/*r*HsDHODH <3), resulting in the selection of 1,948 compounds. The antimalarial activity of these compounds was subsequently tested against *P. falciparum* asexual blood-stage parasites. As a result, 758 compounds exhibited IC_50_ values < 25 µM, yet there was no clear correlation between *r*PfDHODH and antimalarial activity (Fig. S2g), likely due to low membrane permeability or off-target effects among the library compounds. The potential of compounds with antimalarial activity to exhibit toxicity was evaluated using HepG2 cells (Fig. S2h) for general cytotoxicity and isolated porcine mitochondria (Fig. S2i) for electrotransport inhibition. Compounds with a CC_50_ value >25 µM in both assays were selected as hits. To further narrow the hit list, a subset of 730 compounds was clustered based on chemical fingerprints, resulting in 228 clusters. These clusters were subsequently reduced to 168 based on structural novelty and then to 43 clusters of hits exhibiting desirable characteristics. Based on compound purity and the absence of conflicting interests due to intellectual property rights, the 43 clusters (69 compounds) were reduced to 35 clusters including 53 compounds, which were then subjected to cross-inhibition analysis against *r*PvDHODH and *r*PbDHODH using an optimized 2,6-dichlorophenolindophenol (DCIP)-based enzyme assay (Fig. S1). In both cases, the assay system was robust in terms of hit identification, as illustrated by the average Z’ factor of 0.82 and average DSM265 reference control IC_50_ of 130 nM for *r*PbDHODH and average Z’ of 0.72 and DSM265 inhibition IC_50_ of 20 nM for *r*PvDHODH (Fig. S2j-l). The inhibitory activity of DSM265 against both recombinant enzymes was consistent with previously reported values (Coteron et al., 2011; Phillips et al., 2016). This process resulted in a further reduction of the hit list to 348 compounds exhibiting a safe profile (Fig. 1a).

**Figure 1.**
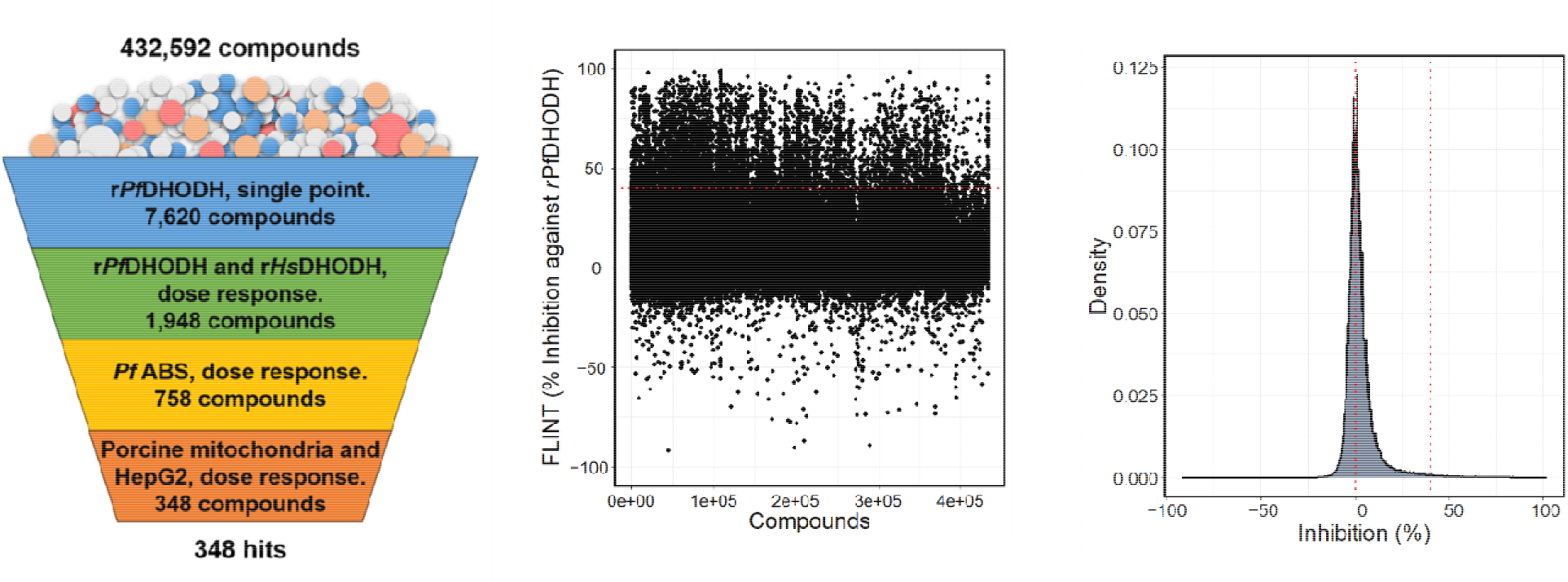
Screening workflow and hit identification criteria. (a) Target-based high-throughput screening assay scheme. (b) Density plot of the dataset at primary screening against PfDHODH, showing the normal distribution of the primary screening data. The red line through zero represents the average inhibition at a single dose. By contrast, the other vertical line represents the cut-off inhibition established by the empirical rule of the normal distribution. (c) Scatterplot of early hit identification with hit selection criteria set to 32% inhibition after a single-dose assay against PfDHODH.

### 2.2 Identification of Compounds Exhibiting Potent PfDHODH Inhibition and High Selectivity

Based on potent inhibition of the *P. falciparum* asexual blood stage and recombinant parasite enzymes, a subset of compounds was selected for co-crystallization with *r*PfDHODH. Seven of these compounds yielded *r*PfDHODH co-crystal structures (Table S3), of which four compounds inhibited all recombinant parasite enzymes except *r*HsDHODH and showed no detectable cytotoxicity toward HepG2 cells or inhibition of porcine mitochondria (Fig. 2 and Table 1).

**Figure 2.**
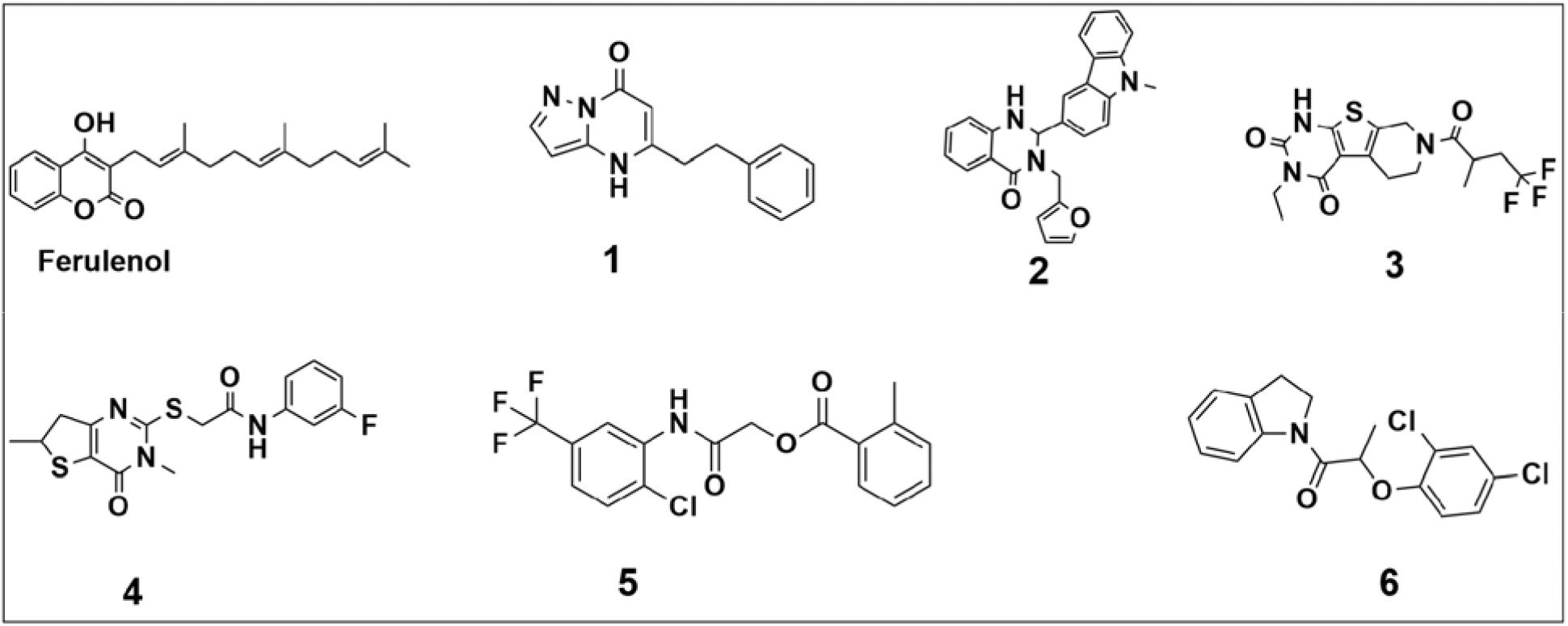
Chemical structures of compounds exhibiting strong inhibition against PfDHODH.

**Table 1.** Selected compounds identified by high-throughput target-based screening and confirmed by whole-cell–based assays against the parasite. **ABS**: asexual blood stage; **EC_50_**: half-maximal concentration required to inhibit parasite viability; **IC_50_**: half-maximal concentration required to inhibit recombinant protein.

| Compounds | PfDHODH<br>IC<sub>50</sub> (μM) | PvDHODH<br>IC<sub>50</sub> (μM) | PbDHODH<br>IC<sub>50</sub> (μM) | EC<sub>50</sub> ABS<br>(μM) | HsDHODH<br>IC<sub>50</sub> (μM) | Selectivity<br>index |
| --- | --- | --- | --- | --- | --- | --- |
| **Ferulenol** | 0.1 | 0.07 | 0.18 | 0.56 | 7.07 | 71 |
| **1** | 0.11 | 4.43 | 6.47 | 4.44 | 11.22 | 102 |
| **2** | 0.06 | 0.01 | 3.59 | 1.21 | 0.22 | 4 |
| **3** | 0.37 | 0.14 | 0.22 | 0.4 | 12.33 | 33 |
| **4** | 0.24 | 1.39 | 0.64 | 1.47 | 10.48 | 44 |
| **5** | 0.18 | 0.03 | 0.02 | 1.57 | 0.75 | 4 |
| **6** | 0.36 | 0.04 | 0.05 | 3.56 | 5.83 | 16 |

All seven compounds inhibited *r*PfDHODH at sub-micromolar concentrations, with compound 2 showing the highest potency (IC_50_ = 60 nM) (Fig. 2 and Table 1). Similar activity was observed against *r*PvDHODH, with all compounds except 1 and 4 showing sub-micromolar inhibition; ferulenol and compounds 2, 5, and 6 exhibited nanomolar potency. All compounds except 1 and 2 also showed consistent sub-micromolar inhibition of rPbDHODH. In cell-based assays, all seven compounds inhibited asexual-stage *P. falciparum*, although cellular EC_50_ values were generally higher than the corresponding *r*PfDHODH IC_50_ values, consistent with potential differences in cellular uptake or target accessibility. Ferulenol and compounds 1 and 3 retained sub-micromolar cellular activity, whereas compounds 2 and 4 showed approximately 20-fold and 6-fold reductions in potency, respectively, relative to their enzymatic activity. Compound 3 exhibited cellular potency comparable to its *r*PfDHODH activity, while ferulenol showed an approximately 6-fold reduction. Against *r*HsDHODH, all compounds except 2, 5, and 6 showed substantially reduced inhibition, consistent with the absence of detectable HepG2 cytotoxicity. Accordingly, compounds 1, 3, 4, and ferulenol displayed strong selectivity for *r*PfDHODH, with selectivity indices >30 (Table 1). None of the seven compounds inhibited mammalian mitochondrial complex II–III or complex I–III at the concentrations tested (Fig. S2g–i), supporting their selectivity for parasite DHODH over mammalian mitochondrial respiratory complexes.

### 2.3 X-ray crystal structures of selected compounds in the PfDHODH active site

X-ray crystallographic data and refinement statistics are summarized in Table S2. Ferulenol binds between Ile263 and Phe171, with its coumarin benzene ring occupying a hydrophobic pocket formed by Val532, Ile263, and Phe227. The pyran ring extends toward the adjacent polar region, where its α-pyrone carbonyl and ring oxygen interact with Arg265. The carbonyl oxygen also forms hydrogen bonds with a water molecule and Cys184, while the phenolic hydroxyl group directly hydrogen bonds with His185. Its prenylated side chain extends toward Gly535 and the hydrophobic region near Cys175 and Phe171, further stabilizing the ligand (Fig. 3a). Compound 1 is anchored by hydrogen bonds between its pyrimidine moiety and Arg265, Cys184, and His185, while its benzene side chain occupies a hydrophobic pocket formed by Phe188, Leu197, and Met536 (Fig. 3b). Compound 2, the most potent PfDHODH inhibitor (IC = 0.06 μM), was resolved at 2.55 Å. Six PfDHODH–compound 2 complexes were present in the asymmetric unit, with chain A showing the clearest ligand density. The dihydroquinazoline core is positioned between His185 and Arg265, with His185 hydrogen bonding to the ligand ketone oxygen (O4). The carbazolyl substituent extends into a hydrophobic pocket lined by Tyr168, Phe171, Leu172, Cys175, Cys184, Leu191, and Met536, while the furylmethyl group forms a π–π interaction with Phe188 (Fig. 3b). Compound 3 (IC_50_ = 0.37 μM) forms hydrogen-bonding interactions with Tyr528, His185, and Arg265, with its fused heteroaromatic core accommodated within the active site. Its trifluoroalkyl group extends into the adjacent hydrophobic region near Leu172, Met536, and Pro188 (Fig. 3c). Compound 4 is predominantly stabilized by hydrophobic interactions at both ends of the molecule. Its fluorophenyl group occupies a pocket near Gly181 and Ile263, while the central acetamide forms hydrogen bonds with Cys184 and His185. The thienopyrimidin-4-one moiety extends toward a second hydrophobic pocket, adopting an orientation similar to that of ferulenol (Fig. 3d). The predominantly hydrophobic binding modes of compound 4 and compounds 1, 2, 5, and 6 may contribute to their relatively modest in vitro potency (Table 1).

**Figure 3.**
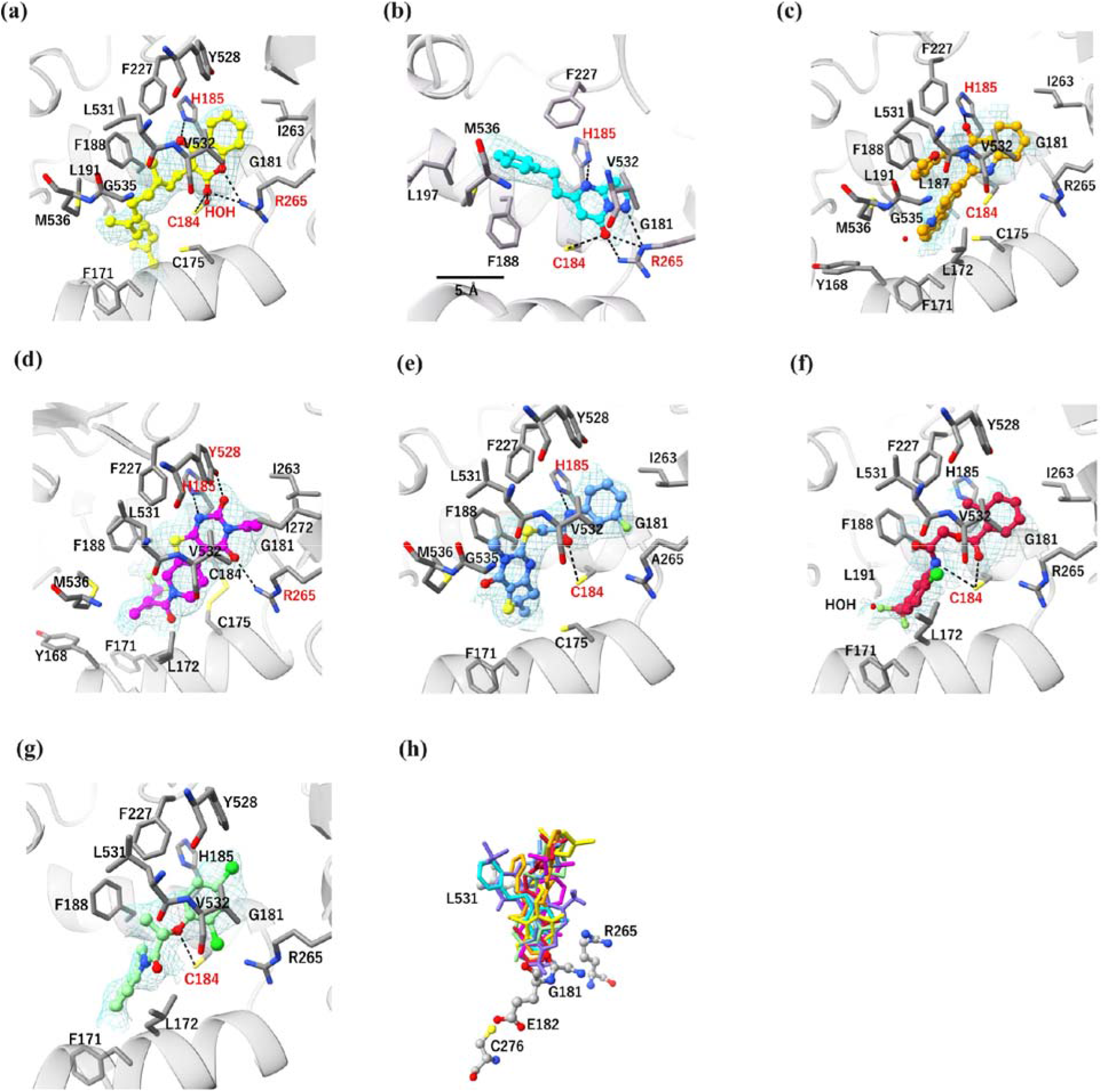
Electron density maps of PfDHODH inhibitors and interaction modes. The omit (*F*_O_-*F*_C_) electron density maps are shown as blue meshes, and hydrogen bonds are indicated by black dashed lines. Protein residues within 4 Å of each ligand are shown as gray sticks. (a) Ferulenol, contoured at 2.0σ and 1.0σ; (b) compound 1 (cyan); (c) compound 2 (orange); (d) compound 3 (magenta); (e) compound 4 (purple); (f) compound 5 (red); and (g) compound 6 (green). (h) Superposition of DSM265 (purple) with the selected compounds. The location of the DSM265 resistance mutation site is indicated by a gray ball-and-stick representation.

### 2.4 Structural orientation of the compounds in the hydrophobic pocket of the active site accounts for selectivity for PfDHODH and potency

All PfDHODH-compound complexes were superposed onto the HsDHODH-brequinar (PDB ID: 1D3G) and PfDHODH-DSM265 (PDB ID: 5BOO) structures to investigate the structural basis of parasite selectivity. Brequinar is a potent biphenyl quinoline carboxylic acid inhibitor of HsDHODH (IC_50_ = 0.48 nM) and was the first DHODH inhibitor to enter clinical evaluation for solid tumors (DeRatt et al., 2020). In general, all compounds exhibited convergence of the aromatic head groups within the binding pocket of both DHODH orthologs. However, substantial differences were observed in the orientation of the remaining portions of the molecules along the HsDHODH binding tunnel, resulting in three distinct conformational patterns (Fig. 4). Generally, ferulenol and compounds 3-6 adopted a curved conformation, in which the tail groups were directed toward one side of the active site. By contrast, brequinar maintained a relatively linear conformation and occupied the central region of the tunnel. Importantly, compounds 1 and 2 adopted distinct conformations, particularly with regard to the aromatic side-chain moieties, which were oriented in opposite directions relative to ferulenol and brequinar (Figs. 4 and S4a). Despite its relatively low molecular weight (239.3 Da), the benzene ring of compound 1 was positioned in a manner that generates steric hindrance within the hydrophobic pocket of HsDHODH. By contrast, compound 2 appears to fit well into the open hydrophobic tunnel of the active site, where the furylmethyl group is affected by minimal steric interference, aligning with the binding mode of brequinar in the active site. This structural difference likely accounts for the high selectivity of compound 1 and the reduced selectivity of compound 2 (Fig. S4a). Notably, both compounds exhibit a similar binding conformation in the aromatic head groups, emphasizing that selectivity is primarily modulated by interactions within the hydrophobic binding pocket of DHODH rather than the conserved head-group interactions (Fig. S4a).

**Figure 4.**
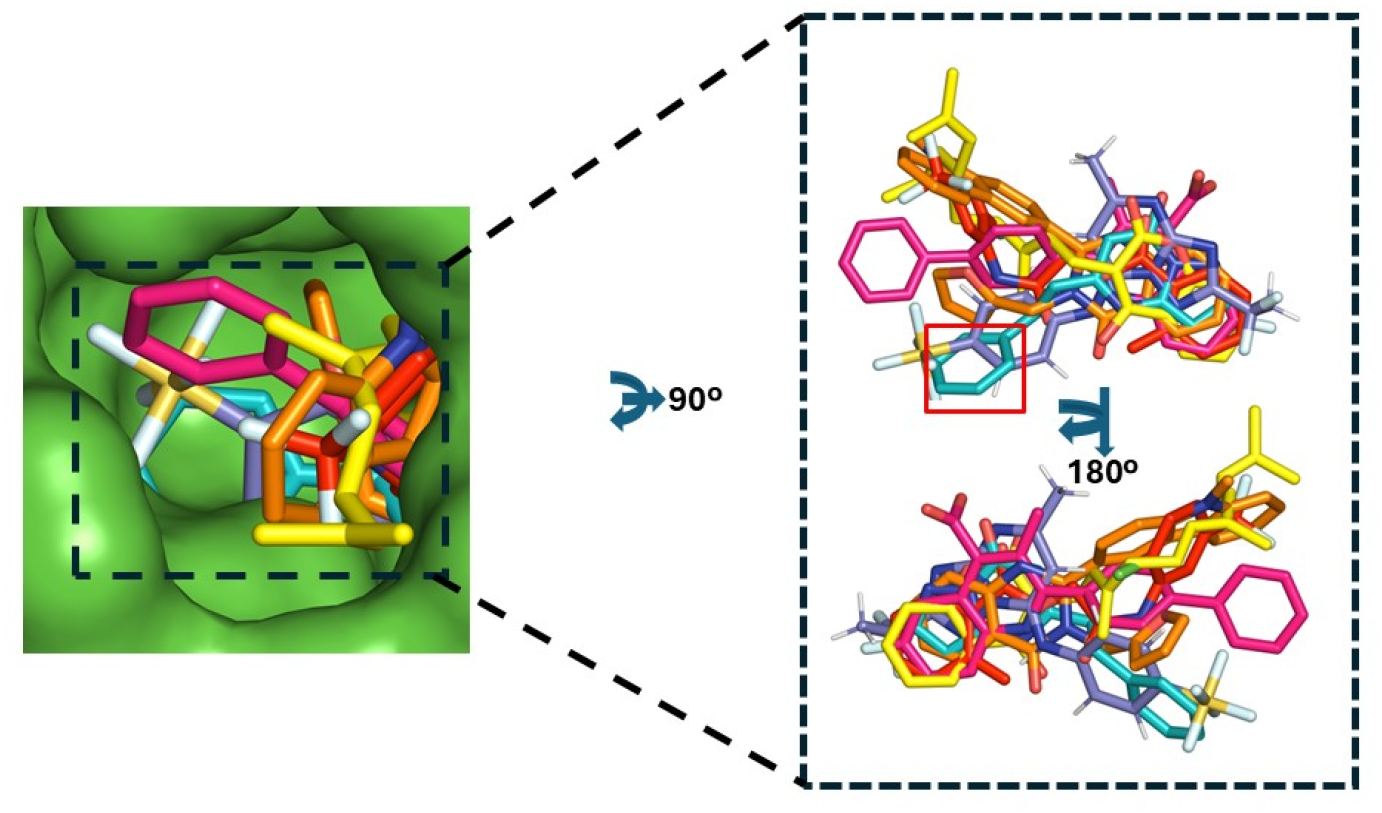
Structural comparison of PfDHODH inhibitors with the HsDHODH inhibitor-binding site. Surface representation of HsDHODH (green) overlaid with the crystal structures of PfDHODH in complex with DSM265, compounds 1 and 2, and ferulenol. The HsDHODH-brequinar complex is shown in pink. Inhibitors are displayed as stick representations, and the colors are consistent with those used in Figure 3. The superposition illustrates the relative orientations of the inhibitors within the human DHODH binding site and facilitates comparison of features associated with host enzyme inhibition.

Despite its strong enzymatic inhibition and high selectivity, the comparatively weaker potency of compound 1 in whole-cell parasite assays may reflect the compound’s higher lipophilicity, which could inhibit cellular uptake or intracellular bioavailability. We previously reported the co-crystal structure of ferulenol and human DHODH (Sato et al., 2020). In HsDHODH, the ferulenol coumarin core is flipped relative to its orientation in PfDHODH, resulting in a corresponding repositioning of the prenylated chain (Fig. S4a). Consequently, the aromatic scaffold undergoes a slight adjustment, whereas the prenylated chain bends in the opposite direction toward a pocket formed by Phe98 and Leu359, the latter corresponding to Phe188 and Met536 in PfDHODH, respectively (Figs. 4 and S4a). These differences highlight the conformational plasticity of ferulenol and suggest that species-specific active-site architectures affect ligand binding modes.

These structural differences provide a mechanistic basis for the 100-fold and 70-fold selectivity of compound 1 and ferulenol for the parasite enzyme, respectively (Table 1). These findings are consistent with the well-established observation that the most significant structural differences between the human and parasite active sites occur within the hydrophobic binding pocket, underscoring its central role in determining selectivity (Pippione et al., 2026). The high potency of compound 2 is likely driven by additional hydrophobic interactions mediated by its furylmethyl group. However, its sterically unhindered orientation within the hydrophobic tunnel of HsDHODH closely resembles that of brequinar, which likely explains the reduced selectivity of compound 2 for PfDHODH over HsDHODH. A similar binding mode was observed for compound 5, the fluoromethyl group of which also occupies the open hydrophobic tunnel without steric interference, consistent with its diminished selectivity. A trend similar to that of ferulenol and compound 5 was observed for compounds 3, 4, and 6 (Fig. S4a). Considered together, the current co-crystal structures provide a structural framework for the rational optimization of these compounds to improve parasite selectivity.

### 2.5 Comparison of selected compounds with DSM265 reveals opportunities for improving inhibitor efficacy

We next compared the binding modes of the selected compounds with the crystal structure of DSM265, the most clinically advanced PfDHODH inhibitor. DSM265 occupies a binding position between His185 and Arg265, like all of the compounds examined in this study (Figs. 3h and S4c). We mapped the binding orientations of the selected compounds relative to residues associated with resistance mutations observed in PfDHODH under DSM265 pressure *in vitro*, including Gly181, Glu182, Arg265, Cys276, and Leu531. In PfDHODH, DSM265 overlaps with compound 1 in the binding site but not with those of ferulenol or compounds 2-6, suggesting that these compounds may retain activity against DSM265-resistant strains (Fig. 3h).

The binding orientation of DSM265 (purple in Fig. 3h) appears to closely overlap with the binding orientations of the selected compounds, with only minor differences in the positioning of specific aromatic substituents. With ferulenol, the principal distinction lies in the orientation of the isoprenyl moiety compared with the DSM265 conformation. Likewise, ferulenol largely superimposes with compound 1, differing primarily in the methyl-carbazol substituent. In both ferulenol and compound 1, the isoprenyl tail and 9-methyl-9H-carbazol moiety, respectively, extend towards the hydrophobic pocket of PfDHODH (Fig. 4). Occupation of the hydrophobic pocket is a key determinant of PfDHODH inhibitor potency, as demonstrated by DSM265. Although the terminal phenyl ring of compound 1 and the furylmethyl group of compound 2 occupy similar regions of the binding site, the latter adopts a conformation that enables the formation of more extensive hydrophobic contacts with surrounding residues (Fig. 3h). This difference likely underlies the marked difference in efficacy between compounds 1 and 2 (Table 1). Collectively, these findings identify key structural regions for future structure-activity relationship (SAR)-guided optimization of compound potency and resistance profiles.

## 3. Discussion

Malaria remains a major global health challenge, and the emergence of resistance to frontline antimalarial drugs continues to drive the need for new therapeutic agents (World Malaria Report 2025; Okwu et al., 2025). PfDHODH is among the most extensively validated antimalarial targets and has demonstrated potential for both treatment and chemoprevention, with activity against blood-and liver-stage parasites (Deng et al., 2009; Nie et al., 2025). Although the PfDHODH inhibitor DSM265 progressed to clinical development, its associated liabilities highlight the continued need for chemically diverse inhibitors with improved drug-like properties (Phillips et al., 2016). Here, screening of more than 400,000 compounds against recombinant PfDHODH identified seven inhibitors for which co-crystal structures were obtained, providing a structural basis for understanding their binding modes and selectivity.

Structurally, PfDHODH adopts a β/α-barrel fold, with inhibitors binding within a pocket located between N-terminal α-helix 1 (residues 162-176), N-terminal α-helix 2 (residues 181-194), and the core of the barrel domain (Fig. S3). This site lies adjacent to the FMN cofactor and is largely hydrophobic in nature. The site is defined by two key characteristics: a hydrogen-bonding region involving Arg265 and His185, and an adjacent hydrophobic pocket lined by residues contributed, in part, by helices 1 and 2. This ability to hold two pockets makes the protein extremely versatile, as it allows binding to multiple scaffolds and interactions with different classes of inhibitors. Sequence differences between PfDHODH and HsDHODH within this region are considered important contributors to parasite selectivity (Lima Costa et al., 2023).

Consistent with previous biochemical and structural studies, our compounds primarily engaged the conserved His185 and Arg265 residues, which are known to play important roles in inhibitor recognition and sensitivity (Heikkilä et al., 2007).

Over the past two decades, several chemically distinct PfDHODH inhibitor classes have been reported, including phenylbenzamides, naphthamides, triazolopyrimidines, isoxazolopyrimidines, and pyrrole derivatives (Singh et al., 2017). Among the associated compounds, the triazolopyrimidine derivative DSM265 advanced to clinical evaluation and remains one of the most important benchmarks in PfDHODH-directed antimalarial discovery (Llanos-Cuentas et al., 2018; Pippione et al., 2026). Our structural analyses (Fig. S4b) showed that the substituted carbonyl group at the C2 position of ferulenol forms a direct hydrogen bond with the conserved residue His185 in PfDHODH, whereas an equivalent interaction is absent with the corresponding His56 residue in the human ortholog (Fig. S5). This distinction appears to be an important determinant of inhibitor selectivity toward the parasite enzyme. As previously reported, the analogous interaction in human and rat DHODHs is typically mediated by an ordered water molecule bridging the inhibitor and His56, rather than by a direct hydrogen bond. Because water-mediated interactions are generally weaker than direct hydrogen bonds involving the histidine imidazole group, the direct interaction of PfDHODH with His185 likely contributes to greater binding affinity and thereby enhanced parasite selectivity (Deng et al., 2014). By contrast, compound 2 represents a substituted hydroquinazoline scaffold and displays distinct interaction patterns within the hydrophobic binding region. Unfortunately, the sterically unhindered alignment of compound 2 within the HsDHODH active site, similar to that of compound 5, likely accounts for the reduced selectivity of compound 2 for PfDHODH over HsDHODH. Compounds 1, 3, and 4 (Table 1 and Fig. 2) contain pyrimidine-based scaffolds. However, to our knowledge, this study is the first to describe both the antimalarial activity and corresponding co-crystal structures of these compounds in complex with PfDHODH. Additionally, compounds 3 and 4, together with compounds 5 and 6 (Table 1), formed hydrogen-bonding interactions with Cys184 in PfDHODH, corresponding to Ala55 in HsDHODH (Fig. S4b). Compounds 5 and 6, which exclusively rely on hydrogen-bonding interactions with Cys184 in PfDHODH (corresponding to Ala55 in HsDHODH), also exhibited the lowest selectivity indices for PfDHODH over HsDHODH.

We previously reported the inhibitory activity of ferulenol against DHODH orthologs from *Eimeria tenella* (IC_50_ = 483 nM), HsDHODH (IC_50_ = 4.7 nM), and PfDHODH (Sato et al., 2020). In that study, ferulenol showed no detectable inhibition of recombinant PfDHODH, even at concentrations up to 20 µM. By contrast, the present study revealed substantially stronger activity against recombinant PfDHODH, with an IC_50_ of 0.1 µM. This discrepancy likely reflects differences in assay methodology, particularly the use of a DCIP-based assay in the previous study compared with a resazurin-based assay employed here. In a separate study, we identified ferulenol as a dead-end inhibitor of *P. falciparum* malate oxidoreductase (PfMQO), selectively binding the PfMQO-dUQ complex and blocking product formation independent of the initiating substrate. Although these previous biochemical analyses demonstrated potent inhibition (IC_50_ = 57 nM), the physiological relevance of the proposed binding site remained unresolved (Hartuti et al., 2018). Subsequent reports suggesting that MQO is dispensable for parasite survival further reduced its attractiveness as a primary antimalarial target (Rajaram et al., 2022). The PfDHODH-ferulenol co-crystal structure reported here, by contrast, provides direct structural evidence supporting PfDHODH as a major molecular target of ferulenol within the parasite mitochondrion. Nevertheless, our earlier observations involving PfMQO, together with multiple reports describing the broad biological activities and toxicity profiles, raise the possibility that ferulenol acts as a multitarget inhibitor affecting multiple components of the MtETC (Fraigui et al., 2002; Kabongo et al., 2023; Lahouel et al., 2007; Niramolyanun et al., 2025).

Collectively, our findings provide the first protein-inhibitor X-ray crystal structures for these chemotypes in complex with PfDHODH, thereby establishing a structural framework for future SAR studies aimed at improving both efficacy and safety. Importantly, five of the seven hit compounds presented in Table 1 satisfied previously proposed hit-to-lead criteria for potential antimalarial compounds, including >10-fold selectivity indices and cellular EC_50_ values <1 μM (Katsuno et al., 2015). In addition, these compounds exhibited sub-micromolar inhibitory activity against DHODH orthologs from multiple parasite species, including the rodent malaria parasite *P. berghei*, supporting their potential *in vivo* antimalarial activity, which warrants further investigation.

## 4. Materials and methods

### 4.1 Initial screening against *r*PfDHODH and *r*HsDHODH

For screening of the in-house library of Shionogi & Co., Ltd. compounds, high-throughput screening assays with purified *r*PfDHODH and *r*HsDHODH were established based on a previous method, with optimization (Caballero et al., 2016). An Echo 555 liquid handler (Labcyte, Sunnyvale, CA) was used to dispense 12.5 nL of each compound in DMSO or DMSO alone as a control into a 1536-well black plate (Greiner Bio-One, Monroe, NC), which was then sealed and stored at −30°C until use. For the *r*PfDHODH single-dose assay, a 1 mM compound stock solution was used, whereas for the *r*PfDHODH and *r*HsDHODH dose-response assays, a 4 mM compound stock solution with 9 serial dilutions in 2 steps was used and tested in triplicate. Two microliters of 100 µM L-dihydroorotic acid (L-DHO) and resazurin in assay buffer (100 mM HEPES [pH 7.0], 5 mM CHAPS, 0.05% BSA, 150 mM NaCl, 100 nM FMN) was dispensed into each well of the compound plates using a FlexDrop PLUS (Perkin Elmer, Waltham, MA), followed by the dispensing of 2 µL of 0.6 µg/mL *r*PfDHODH or *r*HsDHODH in assay buffer in the same manner. After a 90-min incubation at 25°C, the enzyme reaction was stopped by adding 4 µL of 10 mM orotate containing 0.005% (w/v) KM-70, and the fluorescence (Ex 540 nm/Em 590 nm) of resorufin was measured using a PHERAstar FSX microplate reader (BMG LABTECH GmbH, Ortenberg, Germany).

### 4.2 Counter screening assay against *r*PvDHODH and *r*PbDHODH

The reaction conditions for analyzing *r*PvDHODH and *r*PbDHODH were first optimized to fit the 96-well plate (BMBiO #P7530728) (Fig. S1 and Table S1) format using a DCIP calorimetric assay. Briefly, 195 µL of assay mixture, consisting of assay buffer (100 mM HEPES [pH 7.6], 5% glycerol, 150 mM NaCl, 0.05% Triton X-100), 120 μM DCIP, 15 μM decylubiquinone, and 25 μg/mL of *r*PvDHODH membrane protein, was dispensed into the wells of a 96-well plate containing 1 μL of test compound per well. The plates were then mixed at 800 rpm using a MixMate Eppendorf plate mixer, and the speed was increased to 1000 rpm for 30 s. Thereafter, 5 μL of substrate (8 mM L-DHO) was added to start the reaction, and the plate was mixed as described above. The absorbance of each well was recorded at 600 nm after 10 min using a SpectraMax Paradigm plate reader. A single concentration of DSM265 (100 μM final) was used as the positive control, and DMSO was used as the negative control. The same assay conditions were repeated for PbDHODH.

### 4.3 Crystallization

For crystallization, purified PfDHODH was replaced with crystallization buffer (10 mM HEPES [pH 7.6], 300 mM NaCl, 5% glycerol, 10 mM DTT, 0.1% [v/v] Tween20) using a centrifugal filter unit (Amicon Ultra-30, 30-kDa MWCO, Millipore). To the crystallized sample, 1 mM L-DHO and possibly 0.5 mM candidate inhibitor compound were added to make a protein solution. Drops were made so that the volume ratio of the protein solution to the mixed solution was 1:1, and crystallization was performed using the sitting-drop vapor diffusion method. Crystallization conditions were initially screened at 293 K using the sitting-drop vapor diffusion method in a 96-well plate (Violamo Protein Crystallization Plate, AS ONE, Osaka, Japan). A 0.5-mL droplet containing approximately 20 mg/mL of PfDHODH dissolved in crystallization buffer (10 mM HEPES [pH 7.6], 300 mM NaCl, 5% glycerol, 10 mM DTT, 1 mM L-DHO supplemented with 0.5 mM inhibitor [ferulenol and compound 2]) was mixed with an equal volume of reservoir solution, and the droplet was allowed to equilibrate against 80 mL of reservoir solution. Other inhibitors (compounds 1, 3-6) were soaked after co-crystallization with 1 mM L-DHO was achieved. The concentration of the inhibitors was 1 mM. After optimization of all conditions, the best crystals were obtained with 0.15-0.21 M sodium acetate [pH 5.9-6.3], 19-20% (w/v) PEG 3400 as the reservoir solution. The best crystals were grown to dimensions of 0.2 × 0.2 × 0.05 mm in 7-14 days at 293 K. The SMILES codes for the selected compounds described in this study are available in Table S3.

## 5. Description of supplementary material

The Supporting Information is provided as a separate file and contains additional methods, graphs, tables, and figures of some of the selected compounds. Atomic coordinates and structure-factor data have been deposited in the Protein Data Bank under accession codes 45DO (PfDHODH with ferulenol), 45EQ (PfDHODH-compound 1), 45FO (PfDHODH-compound 2), 45FP (PfDHODH-compound 3), 45FQ (PfDHODH-compound 4), 45FR (PfDHODH-compound 5), and 45FS (PfDHODH-compound 6). Additional data supporting the findings of this study are available from the corresponding author upon reasonable request.

## 6. Acknowledgements

We thank all staff members of the Beamline BL44XU at SPring-8 for assistance with X-ray diffraction data collection. Synchrotron-radiation experiments were performed at SPring-8 BL44XU (proposal nos. 2020B6518, 2021A6616, 2021B6616, 2022A6714, 2022B6714, 2023A6814, 2023B6814, 2024A6913, 2024B6913) under the Cooperative Research Program of the Institute for Protein Research, Osaka University. We thank the Japanese Red Cross Society for providing human red blood cells (registry no. R030038).

## 7. Author Contributions

**Japheth Kibet Ng’etich**: Investigation; data curation; formal analysis; writing – original draft; visualization. **Ryuta Ishii**: Investigation; data curation; formal analysis; writing – original draft; visualization. **Hina Yuasa**: Investigation; formal analysis; writing – original draft. **Ryo Hatanaka**: Investigation; formal analysis. **Aiko Hattori**: Investigation; formal analysis. **Jun-ichi Kishikawa**: Investigation; formal analysis; writing – review and editing. **Endah Dwi Hartuti:** Investigation; data curation; formal analysis. **Normalita Eka Pravitasari:** Investigation; data curation; formal analysis. **Kiyoshi Kita**: Conceptualization; writing – review and editing. **Yusuke Tateno**: Conceptualization; resources; writing – review and editing. **Teruhisa Kato**: Conceptualization; resources; writing review and editing. **Takaya Sakura**: Conceptualization; writing – review and editing; supervision; project administration. **Tomoo Shiba**: Conceptualization; formal analysis; data curation; writing – review and editing; funding acquisition; supervision; visualization; project administration; resources. **Daniel Ken Inaoka**: Conceptualization; investigation; data curation; formal analysis; writing – review and editing; resources; funding acquisition; supervision; visualization; project administration.

## 9. Data availability statement

Additional supporting information will be provided with the manuscript. Further information and data supporting the findings of this study are available from the corresponding author upon reasonable request.

## Abbreviations and Symbols

COVID-19: Coronavirus disease 2019
DHODH: dihydroorotate dehydrogenase
LDAO: *N*,*N*-dimethyldodecan-1-amine oxid
L-DHO: L-dihydroorotic acid
ACT: artemisinin combination therapy
MtETC: mitochondrial electron transport chain
FMN: flavin mononucleotide
PfDHODH: Plasmodium falciparum DHODH
PvDHODH: Plasmodium vivax DHODH
PbDHODH: Plasmodium barghei DHODH
HsDHODH: *Homo sapiens* DHODH
MQO: malate:quinone oxidoreductase
SAR: structure-activity relationship
CC_50_: 50% cytotoxic concentration

