## Supplementary Material for "Structural insights and mechanistic inhibition of *Plasmodium falciparum* dihydroorotate dehydrogenase by newly identified chemotypes"

#### 1. Methods

##### 1.1 Expression and purification of recombinant PfDHODH, PvDHODH, and PbDHODH

Recombinant PfDHODH, PvDHODH, and PbDHODH were cloned and expressed as described previously (Hartuti et al., 2021; Sato et al., 2020). Briefly, codon-optimized PvDHODH (PVX\_113330) and PbDHODH (PBANKA\_0102100) were subcloned into pET101 with C-terminal His<sub>10</sub> tags and expressed in *Escherichia coli* BL21 Star™ (DE3). Positive clones were sequence-verified and stored as glycerol stocks at −80°C. PfDHODH comprising residues 158–569 was expressed from pETSUMO/PfDHODH as an N-terminal His<sub>6</sub>-SUMO fusion in *E. coli* BL21 Star™ (DE3). Cells were grown in Terrific Broth, induced with 25 μM IPTG at an OD<sub>600</sub> of 0.4–0.7, and cultured for 16 h at 20°C. PfDHODH was purified at 4°C by cell disruption, clarification, and Ni-NTA affinity chromatography. The protein was washed and eluted using buffers containing C11DAO and imidazole. The His<sub>6</sub>-SUMO tag was removed by SUMO protease digestion, followed by a second Ni-NTA step to remove the cleaved tag and protease. The resulting protein was concentrated and stored at −30°C in glycerol until use. Before crystallization, PfDHODH was buffer-exchanged and further purified by size-exclusion chromatography using an ENrich SEC650 column in crystallization buffer (10 mM HEPES, pH 7.6, 300 mM NaCl, 5% glycerol, 10 mM DTT, and 0.1% Tween 20). Peak fractions were concentrated to an A<sub>280</sub> of 11–12, and protein purity was confirmed by 12% SDS-PAGE. Recombinant HsDHODH was purified according to the previously established protocol (Sato et al., 2020).

### **1.2 Assessment of mammalian complex I and III inhibition (complex II–III and complex I–III assays)**

The 348 hit compounds were evaluated for inhibition of mammalian mitochondrial respiratory complexes using cytochrome *c* reduction assays, as previously described with minor modifications (Luo et al., 2008, Joshi et al., 2011). Human or porcine liver mitochondria were incubated with compounds in an eight-point, threefold serial dilution (28.9 μM–13.2 nM) in assay buffer containing 2 mM potassium cyanide and 100 μM cytochrome *c*. For the complex II–III assay, the reaction was initiated with 175 μM succinate and incubated for 60 min at room temperature. For the complex I–III assay, 5.25 mM NADH was added to initiate the reaction, followed by a 30-min incubation. Absorbance was measured at 550 nm using a SpectraMax Paradigm microplate reader. Antimycin A (1 μM) and DMSO served as positive and negative controls, respectively. Assays using human and porcine liver mitochondria were performed under identical conditions.

### **1.3 Assay of cytotoxicity against HepG2 cells**

HepG2 cells (American Type Culture Collection, Manassas, VA) were suspended in DMEM high glucose (Sigma, D5796) supplemented with 10% fetal bovine serum (Hyclone, SH30070.03) and 1% penicillin-streptomycin mixed solution (Nacalai, 26252-94) and seeded in the wells of a 384-well black plate (Greiner) pre-filled with test compound in DMSO at $4 \times 10^3$ cells/well. An Echo 555 liquid handler was used to dispense 125 nL/well of DMSO or test compound at 5.4 mM, along with 7 serial dilutions in 3 steps, in duplicate. DMSO and digitonin were used as controls. After incubating the plate for 48 h at 37°C in a 5% CO<sub>2</sub> atmosphere, 5 μL of resazurin (TCI, cat. no. R0203) was added to each well with the plate maintained on a plate heater, resulting in a final concentration of 80 µg/mL. The plate was then returned to the incubator and kept for an additional 4 h at 37°C and 5% CO<sub>2</sub>. The reaction products in each well were then detected using a PHERAstar FSX (Ex 540/Em 590).

##### 1.4 Crystallization data collection and structure determination

All X-ray diffraction data were collected at 100 K on a beamline BL44XU ($\lambda = 0.90000$ Å; Rayonix MX300-HE CCD detector, Rayonix, Oak Avenue, Evanston, IL, USA) at SPring-8 (Harima, Japan). For X-ray diffraction experiments at 100 K, a crystal mounted on a nylon loop was transferred to a reservoir solution supplemented with 20% (w/v) glycerol and then flash-frozen in a liquid nitrogen stream. Data were processed and scaled using *XDS* (Otwinowski and Minor, 1997). The structure of *P. falciparum* DHODH in the inhibitor form was solved by molecular replacement using the refined protein coordinates of *P. falciparum* DHODH in complex with DMS265 (PDB code 4RX0) without the inhibitor molecule as a search model (Inaoka et al., 2015). The *MOLREP* program (Vagin and Teplyakov, 1997) as implemented within *CCP4* (The CCP4 Suite, 1994) (http://www.ccp4.ac.uk/), was used for molecular replacement. Crystallographic models were refined to a resolution of 2.15-3.0 Å with the novel inhibitor complex using the program *REFMAC5* (Murshudov et al., 1997). Results of statistical analyses of X-ray data and refinement experiments are summarized in Supplementary Table 2. Graphical representations were generated using *ChimeraX-1.12* (Pettersen et al., 2021). In the complex consisting of compounds 2 and 3, the electron density of the compounds was clearly observable, although the R-value was relatively high because the atomic density seemed to suggest the presence of a surfactant and therefore could not be assigned (Table S2)

### 1.5 Antiplasmodial assay–asexual blood stage

Antiplasmodial activity and target engagement were evaluated using parental *P. falciparum* 3D7 and transgenic 3D7-yDHODH parasites expressing yeast DHODH, as previously described (Hartuti et al., 2021). PfDHODH-targeting compounds were expected to inhibit the parental 3D7 strain but not the transgenic 3D7-yDHODH strain. Parasite viability was determined using a fluorescence-based *P. falciparum* lactate dehydrogenase (PfLDH) assay based on NCOU1 reduction by *E. coli* nitroreductase (NTR) to generate fluorescent umbelliferone (Sakura et al., 2024). The parasite was cultured using standard procedures (Trager and Jensen, 1976) in RPMI 1640 supplemented with 25 mM HEPES (pH 7.4), 0.5% (w/v) Albumax II, and 2% human erythrocytes. Synchronized ring-stage parasites were exposed to test compounds at 0.3% parasitemia and 1% hematocrit in 25 μL assay volumes. DMSO and atovaquone/artemisinin (1 μM each) served as negative and positive controls, respectively, while an eight-point atovaquone dilution series was included as a reference on each plate. Following 72 h incubation at 37°C under 90% N<sub>2</sub>, 5% CO<sub>2</sub>, and 5% O<sub>2</sub>, PfLDH activity was measured by addition of the NTR-NCOU1 detection mixture containing lactate, Tris-HCl, acetylpyridine adenine dinucleotide, NCOU1, NTR, and Triton X-100. After 20 min incubation, fluorescence was measured at 360 nm excitation and 465 nm emission using a PHERAstar Plus microplate reader.

### 2. Supplementary Tables

**Table S1.** Protein activity and optimum concentrations for assay: Membrane protein concentrations used in this study and the optimum concentrations used in the assay.

| Membrane Protein | Total Protein Concentration | Optimum concentration used in assay |
| --- | --- | --- |
| PfDHODH | 25 mg/mL | 0.3 µg/mL |
| PvDHODH | 2647 µg/mL | 25 µg/mL |
| PbDHODH | 1971 ug/mL | 25 µg/mL |

**Table S2.** Data collection and refinement.

| Data set | Compound 1 | Compound 2 | Compound 3 |
| --- | --- | --- | --- |
| <u>Data collection</u> |  |  |  |
| Diffraction source | SPring8 BL44XU | SPring8 BL44XU | SPring8 BL44XU |
| Wavelength (Å) | 0.90000 | 0.90000 | 0.90000 |
| Space group | *P*2<sub>1</sub>2<sub>1</sub>2 | *P*2<sub>1</sub>2<sub>1</sub>2 | *P*2<sub>1</sub>2<sub>1</sub>2 |
| *a*, *b*, *c* (Å) | 185.6, 145.9, 130.4 | 187.6, 146.0, 130.5 | 186.9, 146.0, 130.5 |
| Resolution range (Å) | 50.0-3.00 (3.18-3.00) | 50.0-2.55 (2.70-2.55) | 50.0-2.59 (2.75-2.59) |
| Total no. of reflections | 410,460 | 858,141 | 637,512 |
| No. of unique reflections | 70,691 | 116,517 | 111,074 |
| Completeness (%) | 97.9 (98.0) | 99.7 (97.1) | 99.8 (99.0) |
| Multiplicity | 5.8 (5.69) | 7.4 (7.0) | 5.7 (5.8) |
| *I*/$\sigma$(*I*) | 8.9 (1.9) | 8.5 (2.0) | 9.2 (1.9) |
| CC<sub>1/2</sub> | 0.997 (0.852) | 0.993 (0.891) | 0.998 (0.845) |
| *R*<sub>merge</sub> | 0.139 (0.927) | 0.133 (0.968) | 0.125 (1.106) |
| <u>Refinement</u> |  |  |  |
| Resolution (Å) | 3.00 | 2.55 | 2.59 |
| *R*<sub>work</sub>/*R*<sub>free</sub> | 0.235/0.303 | 0.267/0.308 | 0.246/0.304 |
| No. of non-hydrogen atoms |  |  |  |
| Protein | 17,992 | 17,809 | 17,924 |
| Ligand | FMN (186), ORO (66)<br>Compound 1 (108) | FMN (186), ORO (66)<br>Compound 2 (186) | FMN (186), ORO (66)<br>Compound 3 (156) |
| Solvent | 104 | 172 | 63 |
| Average *B*-factors (Å<sup>2</sup>) |  |  |  |
| Protein | 68.8 | 55.4 | 62.6 |
| Ligand | FMN (50.1), ORO (59.4)<br>Compound 1 (70.4) | FMN (40.2), ORO (57.5)<br>Compound 2 (84.5) | FMN (40.1), ORO (42.2)<br>Compound 3 (53.2) |
| Solvent | 40.6 | 38.4 | 38.2 |
| RMSD |  |  |  |
| Bond length (Å) | 0.016 | 0.018 | 0.016 |
| Bond angle (°) | 1.962 | 2.056 | 1.880 |
| Ramachandran plot (%) |  |  |  |
| Favored regions | 84.4 | 87.0 | 87.9 |
| Allowed regions | 15.5 | 12.5 | 11.9 |
| Outers | 0.1 | 0.5 | 0.2 |
| PDB code | 45EQ | 45FO | 45FP |

| Data set | Compound 4 | Compound 5 | Compound 6 | Ferulenol |
| --- | --- | --- | --- | --- |
| <u>Data collection</u> |  |  |  |  |
| Diffraction source | SPring8 BL44XU | SPring8 BL44XU | SPring8 BL44XU | SPring8 BL44XU |
| Wavelength (Å) | 0.90000 | 0.90000 | 0.90000 | 0.90000 |
| Space group | *P*2<sub>1</sub>2<sub>1</sub>2 | *P*2<sub>1</sub>2<sub>1</sub>2 | *P*2<sub>1</sub>2<sub>1</sub>2 | *P*2<sub>1</sub>2<sub>1</sub>2 |
| *a*, *b*, *c* (Å) | 187.6, 145.8, 130.6 | 186.3, 145.0, 130.2 | 186.8, 145.5, 130.2 | 185.2, 144.6, 130.4 |
| Resolution range (Å) | 50.0-2.40 (2.54-2.40) | 50.0-2.50 (2.65-2.50) | 50.0-2.50 (2.65-2.50) | 50.0-2.15 (2.28-2.15) |
| Total No. of reflections | 843,104 | 642,339 | 733,090 | 1,318,137 |
| No. of unique reflections | 140,560 | 121,029 | 121,932 | 189,784 |
| Completeness (%) | 99.9 (99.8) | 99.4 (98.1) | 99.0 (96.6) | 99.9 (99.4) |
| Multiplicity | 6.0 (5.9) | 5.3 (5.2) | 5.9 (5.4) | 6.9 (7.0) |
| *I*/σ(*I*) | 6.3 (1.5) | 8.9 (2.1) | 8.8 (1.5) | 11.0 (2.0) |
| CC<sub>1/2</sub> | 0.994 (0.850) | 0.995 (0.662) | 0.995 (0.726) | 0.998 (0.876) |
| *R*<sub>merge</sub> | 0.142 (1.048) | 0.134 (1.191) | 0.135 (1.589) | 0.096 (1.013) |
| <u>Refinement</u> |  |  |  |  |
| Resolution (Å) | 2.40 | 2.50 | 2.50 | 2.15 |
| *R*<sub>work</sub>/*R*<sub>free</sub> | 0.232/0.271 | 0.232/0.271 | 0.248/0.309 | 0.226/0.270 |
| No. of non-hydrogen atoms |  |  |  |  |
| Protein | 17,988 | 18,002 | 18,075 | 17,898 |
|  | FMN (186), ORO (66) | FMN (186), ORO (66) | FMN (186), ORO (66) | FMN (186), ORO (66) |
| Ligand | Compound 4 (144) | Compound 5 (150) | Compound 6 (132) | Ferulenol (162) |
| Solvent | 64 | 87 | 98 | 531 |
| Average *B*-factors (Å<sup>2</sup>) |  |  |  |  |
| Protein | 53.2 | 55.2 | 65.1 | 48.9 |
|  | FMN (35.8), ORO (43.5) | FMN (37.9), ORO (41.7) | FMN (46.0), ORO (50.2) | FMN (38.1), ORO |
| Ligand | Compound 4 (69.5) | Compound 5 (121.5) | Compound 6 (120.2) | (41.3) |
|  |  |  |  | Ferulenol (64.0) |
| Solvent | 38.5 | 39.4 | 48.9 | 49.9 |
| RMSD |  |  |  |  |
| Bond length (Å) | 0.010 | 0.010 | 0.011 | 0.006 |
| Bond angle (°) | 1.449 | 1.464 | 1.572 | 1.631 |
| Ramachandran plot (%) |  |  |  |  |
| Favored regions | 90.0 | 90.3 | 83.3 | 91.4 |
| Allowed regions | 9.8 | 9.4 | 10.4 | 8.6 |
| Outers | 0.2 | 0.3 | 0.3 | 0.0 |
| PDB code | 45FQ | 45FR | 45FS | 45DO |

**Table S3.** SMILES codes for the selected compounds described in this study.

| Compound | SMILES code |
| --- | --- |
| **Ferulenol** | CC(=CCC/C(=C/CC/C(=C/CC1=C(C2=CC=CC=C2OC1=O)O)/C)/C)C |
| **1** | O=C1N2C(NC(CCC3=CC=CC=C3)=C1)=CC=N2 |
| **2** | O=C1N(CC2=CC=CO2)C(C3=CC=C4C(C5=C(C=CC=C5)N4C)=C3)NC6=CC=CC=C61 |
| **3** | O=C(C(C)CC(F)(F)F)N1CCC(C(C(N(CC)C(N2)=O)=O)=C2S3)=C3C1 |
| **4** | FC1=CC=CC(NC(CSC2=NC(CC(C)S3)=C3C(N2C)=O)=O)=C1 |
| **5** | O=C(COC(C1=CC=CC=C1C)=O)NC2=CC(C(F)(F)F)=CC=C2Cl |
| **6** | O=C(C(OC1=CC=C(Cl)C=C1Cl)C)N(CC2)C3=C2C=CC=C3 |

#### 3. Supplementary Figures

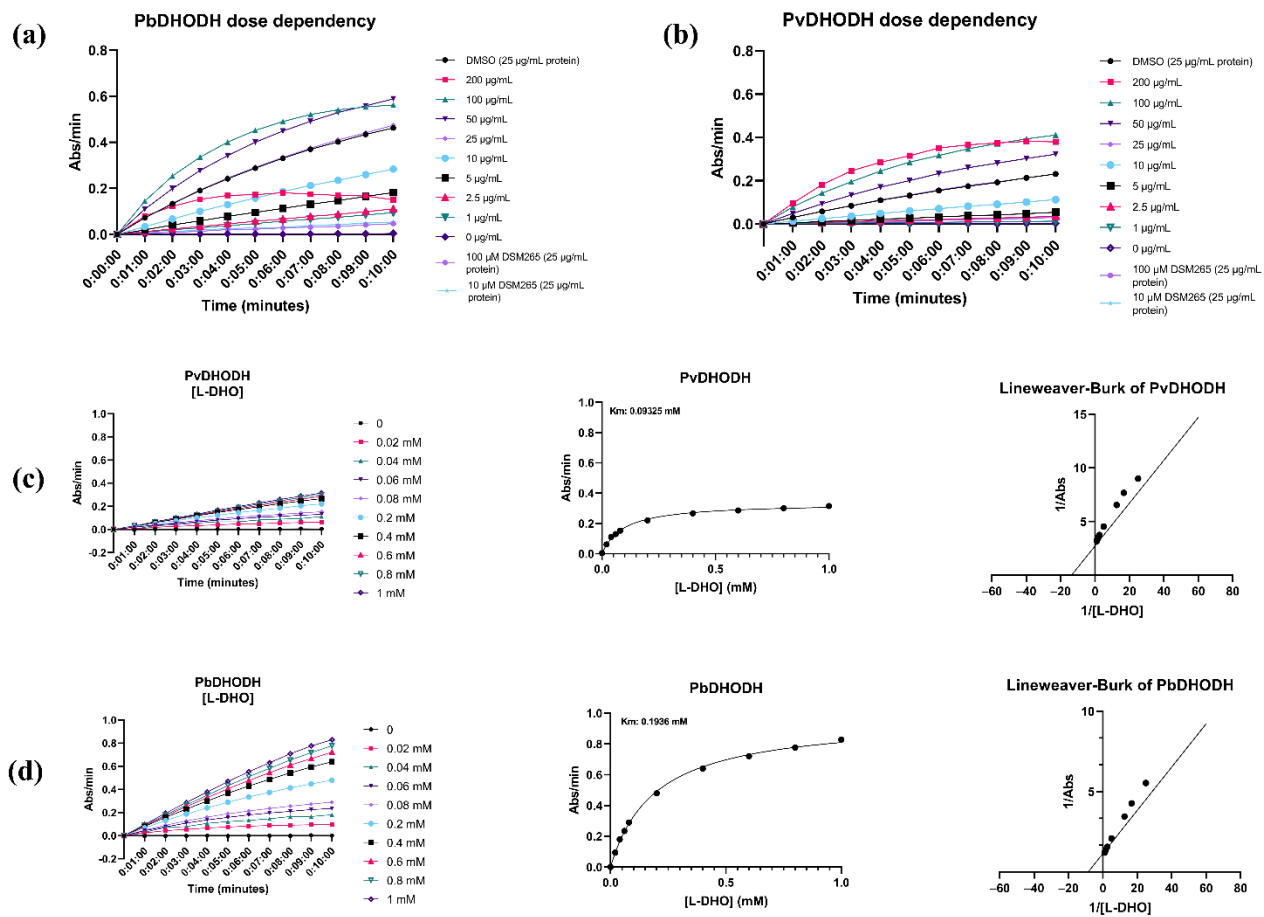

**Figure S1. Optimization of assay conditions for PvDHODH and PbDHODH.** Protein concentration optimization using dose-response assays for PbDHODH (a) and PvDHODH (b), respectively, identifying 25 µg/mL as the optimal enzyme concentration for inhibitor screening. (c, d) Substrate concentration optimization for PvDHODH and PbDHODH, respectively, establishing 200 µM as the optimal substrate concentration for subsequent enzyme inhibition assays.

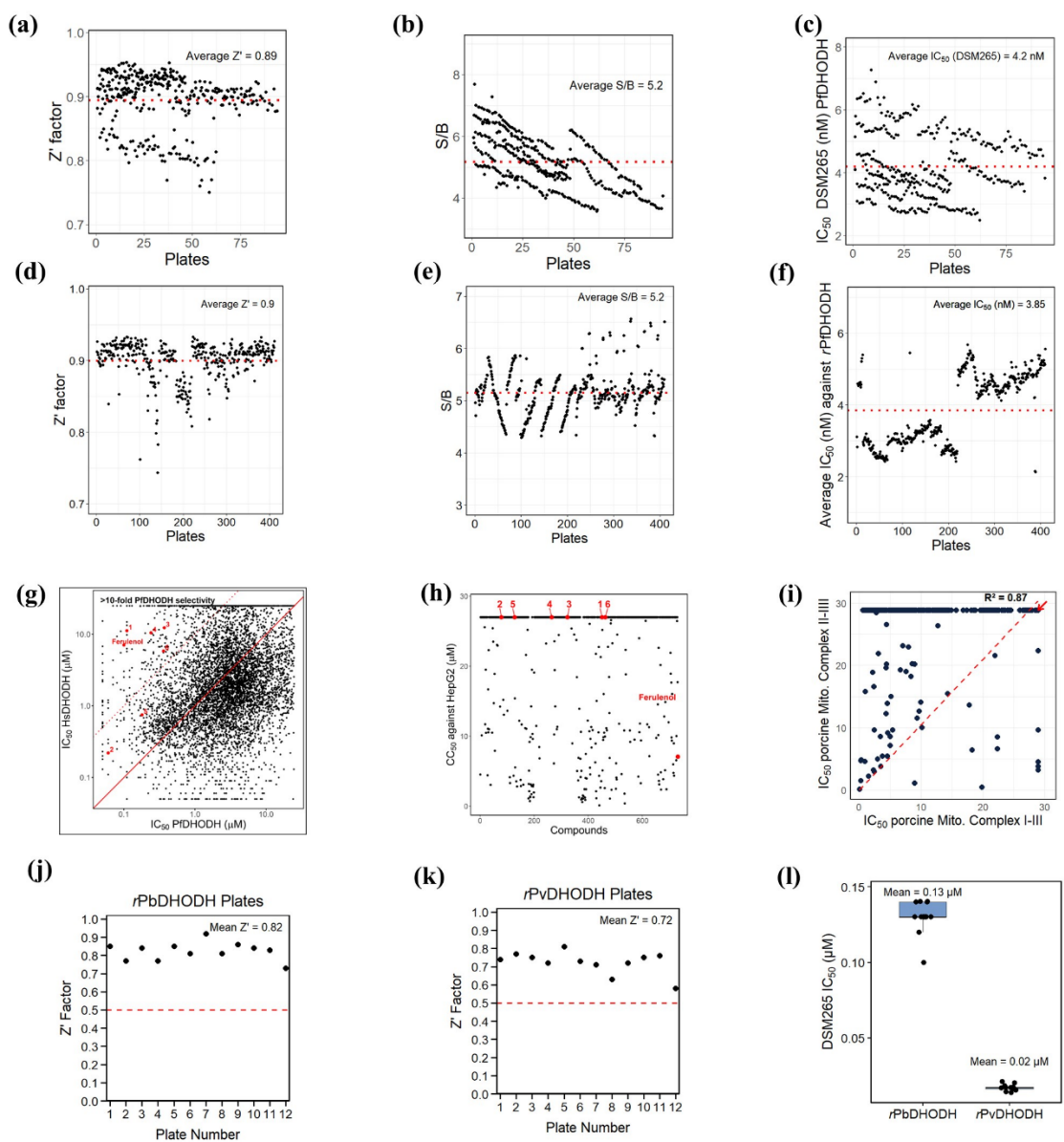

**Figure S2. Assay quality parameters and comparison of compound inhibition profiles across different proteins and assays.** (a) The $Z'$ factor, (b) signal-to-background ratio (S/B), and (c) the $IC_{50}$ of DSM265 against *r*PfDHODH across all assay plates in the first screening. (d) The $Z'$ factor, (e) S/B, and (f) the $IC_{50}$ of DSM265 against *r*PfDHODH, respectively, for dose-response plates. (g) Correlation plot of hit compounds against *r*HDHODH and *r*PfDHODH to assess selectivity for the parasite protein. Compounds exhibiting a $>10$-fold shift in $IC_{50}$ when tested against the human ortholog versus the parasite protein (*r*PfDHODH) were considered

selective for *rPfDHODH* and then selected for further assay. (h) Distribution of $CC_{50}$ values of compounds when tested against the HepG2 cell line. Selected compounds annotated in red showed no cytotoxicity against HepG2 cells, even at the highest tested dose. (i) Assessment of porcine mitochondrial complex I-III and complex II-III for selected compounds. Red arrows show the position of all seven selected compounds described in this study, indicating no inhibition of mitochondrial enzymes. Assay quality parameters of plates used in the selection process for compounds targeting *rPbDHODH* and *rPvDHODH*, including $Z'$ factors (j, k, and l) and $IC_{50}$ values of the reference drug (DSM265).

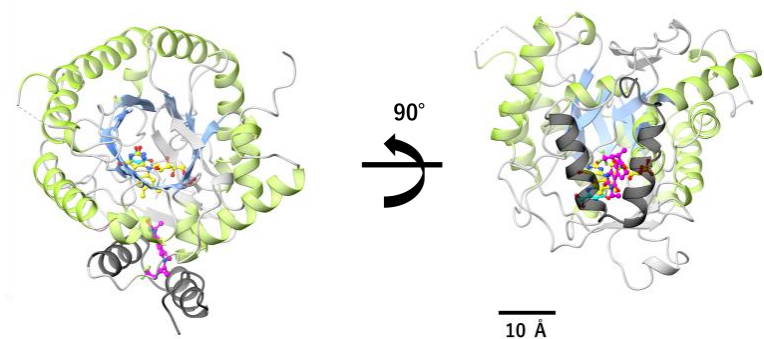

**Figure S3. General structure of PfDHODH with bound compounds in the active site.** The $\alpha/\beta$ TIM-barrel domain is highlighted with α-helices in green and β-strands in blue. The N-terminal microdomain is gray. FMN (yellow), orotate (cyan), and compound 3 (magenta) are bound.

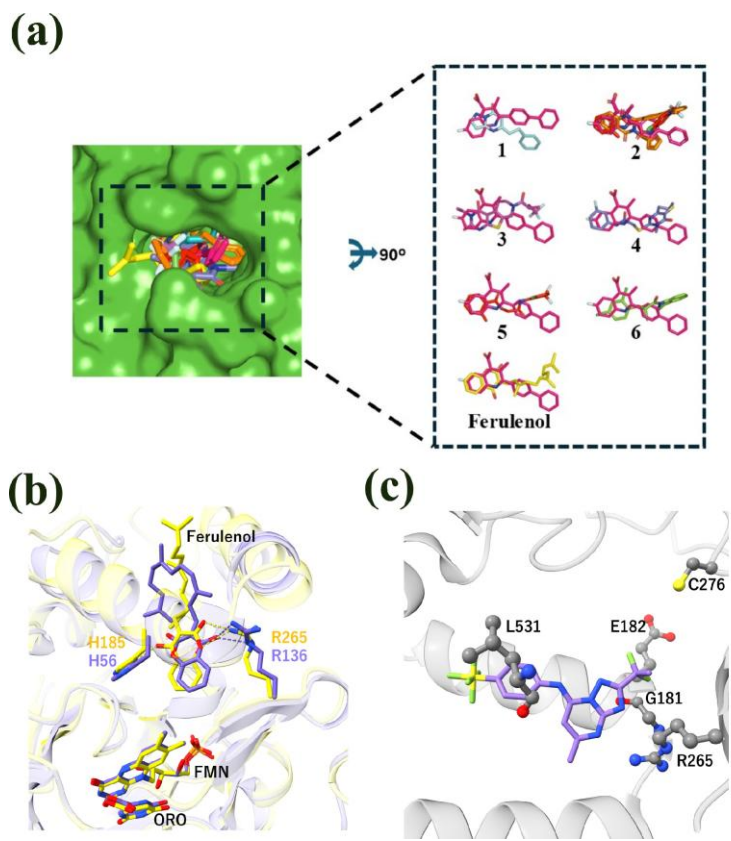

**Figure S4. Superimposed structures showing the orientation of each selected compound relative to the HsDHODH-brequinar complex.** (a) Surface and individual compounds displayed as sticks, with colors consistent with those used throughout the text, compared with the binding orientation of brequinar (pink) within the HsDHODH active site. (b) Detailed binding mode of PfdHODH and DSM265 (shown in medium purple). Residues within 4 Å of the compound are shown as sticks, colored gray. Amino acid residues hydrogen-bonded to the compound are labeled in red, and hydrogen bonds are indicated by black dotted lines. (c) DSM265 in PfdHODH is shown in medium purple, and reported point mutations as gray ball-and-stick models.

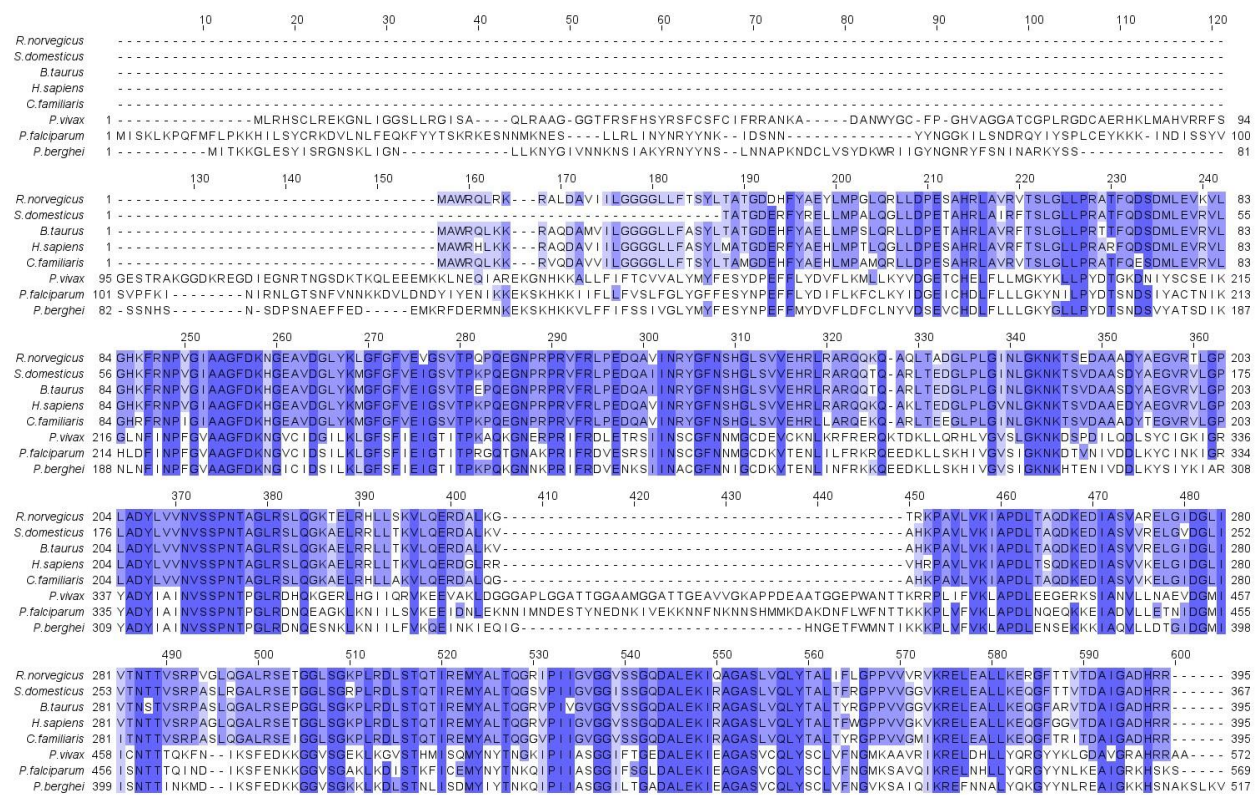

**Figure S5. Multiple sequence alignment of DHODH proteins from selected plasmodial and mammalian species.** Conserved residues are highlighted in dark blue, whereas partially conserved residues are shown in light blue. Residue numbering is based on PfDHODH. Protein sequences were obtained from the National Center for Biotechnology Information Protein database: *P. falciparum* (AAC37170.1), *P. vivax* (KMZ85807.1), *P. berghei* (SCO58773.1), *H. sapiens* (AAH65245.1), *Sus scrofa domesticus* (domestic pig, ALI95909.1), *Canis lupus familiaris* (domestic dog, XP\_038523105.1), *Rattus norvegicus* (rat, CAA56765.1), and *Bos taurus* (domestic cattle, AAI20338.1). Sequence alignment was performed using the European Bioinformatics Institute Multiple Sequence Alignment web server and visualized using Jalview, version 2.11.5.1.
